# Mapping object space dimensions: new insights from temporal dynamics

**DOI:** 10.1101/2024.11.19.624410

**Authors:** A. Kidder, G. L. Quek, T. Grootswagers

**Author notes:** Corresponding author. Department of Psychological and Brain Sciences, Dartmouth College, Hanover, NH, 03755, USA. Laboratory of Brain and Cognition, National Institutes of Mental Health, National Institutes of Health, Bethesda, MD, 20814, USA.

## Abstract

How is object information organized in high-level visual cortex? Recently, a comprehensive model of object space in macaques was proposed, defined by the orthogonal axes of animacy and aspect ratio (Bao et al., 2020). However, when using stimuli that dissociated category, animacy, and aspect ratio in humans, no tuning of aspect ratio was observed in fMRI data (Yargholi & Op de Beeck, 2023). This difference could be a result of different stimuli, the limited temporal resolution of fMRI, or information available about the presented stimuli. Here, we asked if and when information about aspect ratio, animacy, and category is available over time. We collected whole-brain electroencephalography (EEG) data while participants (N = 20) viewed the stimulus set used by Yargholi & Op de Beeck (2023). To mask object details and increase reliance on shape information, we also presented silhouette versions of the stimuli. Stimuli were presented in 5Hz streams using rapid serial visual presentation, with intact and silhouette stimuli sets were shown in separate streams. Using standard multivariate decoding pipelines and representational similarity analysis, we found that aspect ratio, category, and animacy were represented during visual object processing. The dominant dimension was modulated by stimulus type, demonstrating that the observable dimensions of object space depend on the nature of the stimuli presented. Taken together, these findings demonstrate that aspect ratio is represented during object processing, however earlier and more transiently than categorical dimensions, such as animacy. By focusing on underlying temporal dynamics, our results provide clear new insights into the contradicting findings reported in previous work and reveal a more nuanced understanding of how object space evolves over time.

## Introduction

The human visual system is highly skilled at rapidly and accurately recognizing a wide range of visual stimuli. However, the overarching organizational principles used by the visual system to represent these objects is still debated. Several dimensions of object space have been proposed to underlie object space. These dimensions range on a continuum from low level visual statistics (Coggan et al., 2016; O’Toole et al., 2005; Rice et al., 2014) and mid-level properties, such as shape (Bracci & Op De Beeck, 2016; Kayaert et al., 2005; Nasr et al., 2014) and size (Huang et al., 2022; Konkle & Caramazza, 2013; Konkle & Oliva, 2012), to higher level characteristics, such as animacy (Kriegeskorte et al., 2008; Sha et al., 2015), categorical information (Kriegeskorte et al., 2008), humanness (Contini et al., 2020; Grootswagers et al., 2022a), agency (Thorat et al., 2019), and semantic information (Chao et al., 1999; Naselaris et al., 2009). While these diverse features are all represented during object processing, it is still unclear which dimensions primarily drive neural responses, and how these dimensions can best be explained by a unified model of object space.

Recently, a comprehensive map of object space in macaque inferior temporal (IT) cortex was proposed (Bao et al., 2020). This map is characterized by two orthogonal dimensions in which objects can be distributed: the animate-inanimate dimension (animacy) and the stubby-spiky dimension (aspect ratio) (Bao et al., 2020). This map was repeated three times across the hierarchical stages of the ventral visual stream as object representations increased in view invariance. When investigating if this model also exists in humans, representations of and selectivity for all four quadrants of this proposed space (stubby-animate, stubby-inanimate, spiky-animate, spiky-inanimate) were found using 7T functional magnetic resonance imaging (fMRI) (Coggan & Tong, 2023). However, the functional organization found in humans did not mirror the functional organization found in macaques. The stubby-animacy map selectivity highly overlapped with areas traditionally considered selective to different categories (such as face- and body-selective regions), and there were no clear repetitions across hierarchical stages of the visual stream, indicating that while animacy and aspect ratio were fundamental organizing principles, there were differences between the macaque and human visual systems (Coggan & Tong, 2023).

A critique of this map of object space is that the stimuli that were primarily used to identify the aspect ratio and animacy axes did not control for category (Yargholi & Op De Beeck, 2023). In both studies, the stimulus set confounded category and aspect ratio, in that faces were only represented in the stubby-animate quadrant and bodies primarily belonged in the spiky-animate quadrant of object space. When using a new stimulus set that dissociated category from aspect ratio in human participants, no tuning for aspect ratio was found, however category and animacy were strongly represented (Yargholi & Op De Beeck, 2023). Therefore, the authors concluded that aspect ratio does not have a special status for the large-scale organization of object space, and that functional organization of the human ventral visual stream is driven by the dimensions of category and animacy (Yargholi & Op De Beeck, 2023).

While presenting different stimulus sets may be driving these contrasting results, it is also possible that the different neuroimaging techniques utilized by the studies contribute to the seemingly opposing conclusions. In the original macaque study, the stubby-spiky networks were initially found when using electrophysiology, whereas fMRI was used in both human studies (Bao et al., 2020; Coggan & Tong, 2023; Yargholi & Op De Beeck, 2023). It is possible that aspect ratio is used earlier and more transiently than animacy and category information, and due to the slow hemodynamic response, fMRI may not be able to pick up on this dimension as strongly when compared to millisecond-resolution electrophysiology. Additionally, the dimensions that support object processing may be flexible, in that dimensions are weighted differently depending on the object’s available visual information. The macaques viewed “silhouette” objects in conjunction to objects with preserved internal details (Bao et al., 2020), whereas the human participants only viewed objects with preserved internal details (Coggan & Tong, 2023; Yargholi & Op De Beeck, 2023). In conjunction to the varying levels of object detail available in these studies, macaques were shown several objects that they did not have familiarity with (e.g. camera, wheelchair, musical instruments), whereas human participants had category and experiential knowledge about all presented stimuli. Aspect ratio may be used more strongly for objects for which category information is not obvious, or for stimuli that only have silhouette information available, in comparison to known objects that have internal details easily accessible to the visual system.

The goal of the present study was to address these possibilities by using electroencephalography (EEG) and multivariate pattern analysis (MVPA). Using rapid serial visual presentation (RSVP), we presented the stimulus set that dissociated aspect ratio from category, along with the silhouette version of the same stimuli, to human participants in order to directly compare the impact of masking internal features on the object space dimensions used during object processing (Yargholi & Op De Beeck, 2023). Our results indicated that aspect ratio was indeed represented during object processing, however this information was available earlier, more transiently, and less strongly in comparison to animacy and category information for objects with internal details. However, masking object details via silhouettes strengthened aspect ratio information, which became the primary dimension represented, while concurrently weakening animacy and category information. These findings provide insights from a temporal perspective on the functional model underlying object space, and indicate that the dimensions used during object processing is influenced by the visual information we have access to.

## Methods

### Participants

Twenty-two healthy individuals with normal or corrected-to-normal vision volunteered to participate in the study. Participants were recruited from the undergraduate student population and the Western Sydney community. Two participants were excluded due to incomplete recordings, and the data from the remaining twenty participants (13 female, mean age 23.65, SD = 4.04) were included in all analyses. All participants had normal or corrected-to-normal vision. The study was approved by Western Sydney University Institutional Review Board, and informed consent was obtained from all participants prior to the start of the experiment.

### Stimuli

The original stimulus set from Yargholi & Op De Beeck (2023) was used in this experiment, consisting of 52 object stimuli across 4 categories: bodies, faces, manmade objects, natural (Yargholi & Op De Beeck, 2023; Figure 1A). There were 13 stimuli within each category, 6 of which had small aspect ratios and were therefore classified as “stubby,” 6 with large aspect ratios (classified as “spiky”), and 1 that represented the median aspect ratio in that category. Aspect ratio was defined as P^2^/(4**π**A), where P represented the perimeter of the object, and A was the area of the object. All images were grayscale, covered 6.5 x 6.5 degrees of visual angle, and were presented on a light gray background (Yargholi & Op de Beeck, 2023). This stimulus set is referred to as the “intact” stimuli in our paradigm.

**Figure 1.**
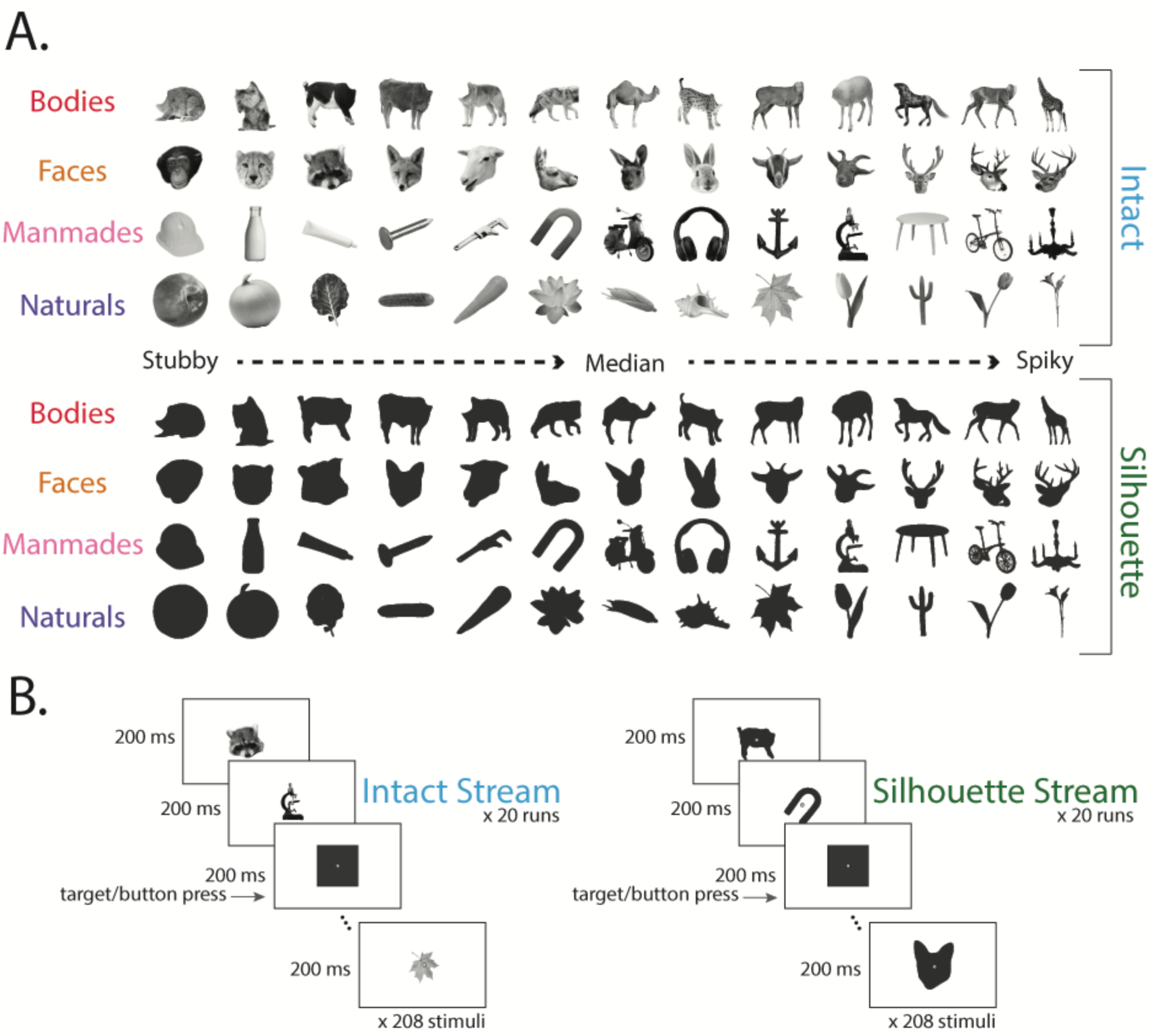
EEG stimuli and experimental design. (a) Visual stimuli used in this experiment included “intact” stimuli (obtained from Yargholi & Op de Beeck, 2023) and silhouette versions of those same stimuli. This stimulus set dissociated animacy from aspect ratio, so that within each category, half of the stimuli were stubby while the other half were spiky. (b) Schematic example of an intact and silhouette stimulus run. Participants fixated on central cross while completing an orthogonal task in which they indicated when a target shape was presented.

Silhouette versions of all stimuli were created to mask internal features of the object stimuli and degrade category object information. Images were binarized using custom code in Matlab. All pixels belonging to the stimulus image were assigned the value of .2 and appeared dark gray. These images were also 256 x 256 and were presented on a gray background. All participants viewed both the intact stimuli and the silhouette version of each stimulus, for a total of 108 stimuli included in the study (Figure 1a).

### Paradigm

Stimuli were presented at a rate of 5 Hz (200 ms/image) in a continuous stream using a rapid serial visual presentation (RSVP) paradigm (Grootswagers, Robinson, & Carlson, 2019; Grootswagers, Robinson, Shatek, et al., 2019; Peirce et al., 2019; Robinson et al., 2019). Each run consisted of a randomized stream of 208 object stimuli (4 repeats of each stimulus), and silhouette images and intact images were always presented in separate runs (Figure 1b). The order the stimuli were presented in were yoked across the intact and silhouette images, so that there were no differences in the presentation order for the different stimulus types. Participants were instructed to fixate on an overlayed center point on the screen, and completed an orthogonal task in which they pressed a button when seeing a target shape of a square or triangle (Figure 1b). There were 2 to 4 target shapes in each run. Runs lasted between 42 - 42.4 seconds, and each participant completed a total of 40 runs of this task (20 runs of the intact stimulus set, and 20 runs of the silhouette stimuli). These yoked runs were presented in a randomized order for all participants. Every stimulus was presented 80 times, for a total of 8,320 (excluding targets) trials during the experiment. The experiment was coded in Python (v 3.6.6.) using the Psychopy library (Peirce et al., 2019). Prior to starting the experiment, participants completed a practice run to familiarize themselves with the speed of stimuli presentation and the task.

### EEG acquisition and preprocessing

Data were continuously recorded using a 64-channel biosemi system at a sampling rate of 2048 Hz. Impedance was reduced below 25 kOhm at each electrode site whenever possible using conductive gel. We used a 64-channel electrode system, arranged according to the international standard 10-10 system. All stimuli were presented on a Viewpixx 120 Hz monitor, and a trigger was sent for the onset of a run, the onset of each stimulus, and the offset of each stimulus.

Minimal preprocessing was performed using the Fieldtrip (version 20231015; (Oostenveld et al., 2011) in MATLAB (version R2022b; The Mathworks, Natick, MA). Data were downsampled to 200 Hz and demeaned to 100 ms prior to stimulus onset (Grootswagers et al., 2017). Epochs of -100 to 800 ms relative to stimulus onset were created for each trial. No additional preprocessing was performed.

### Multivariate Analysis

All following analyses were performed using functions from the CoSMoMVPA toolbox (Oosterhof et al., 2016), and custom code in MATLAB (version R2022b; The Mathworks, Natick, MA). We took a multivariate approach to the data (Grootswagers et al., 2017). To determine if representations of aspect ratio, animacy, and category were present for objects in whole-brain EEG signal, we performed classification analysis. Next, to examine the temporal dynamics and stability of each dimension of object space, we ran a temporal generalization analysis (King & Dehaene, 2014). To test which feature dimension best represented the data across timepoints, we used representational similarity analysis (RSA) and compared model representational dissimilarity matrices (RDMs) to RDMs generated by the EEG data.

### Classification analysis

A regularized linear discriminant analysis (LDA) classifier was trained to decode between whole-brain patterns in the EEG signal across all electrodes at each time point of the trial (Figure 1c). Classifiers were trained to decode between separate conditions (stubby vs spiky, animate vs inanimate, category, individual stimulus) using a leave-one-run-out cross-validation approach. Silhouette stimuli and intact stimuli were analyzed separately from one another. When decoding stubby vs spiky, the object stimuli that were the median for each category were removed from analysis, as they were classified and neither stubby or spiky. The resulting chance-level accuracy was 50% for the classification of animacy and aspect ratio, 25% for stimulus category, and 1.92% for individual stimulus identity (52 images). This classification was performed for each participant individually, and then averaged across participants to obtain the mean classification accuracy across individuals.

### Temporal Generalization Analysis

We performed a temporal generalization analysis to determine the stability of the representations of different dimensions of object space (Carlson et al., 2011; King & Dehaene, 2014). This analysis tested if the representational structure of each dimension was consistent across time points, or if there were shifts in this representation over the time course of object perception. LDA classifiers were trained to discriminate between conditions at each time point and then were tested at all other time points.

### Representational Similarity Analysis

To find the feature model that best fit the EEG data at each time point, Representational Similarity Analysis (RSA) was implemented (Kriegeskorte, 2008). First, to characterize the similarity between each stimulus within whole-brain EEG responses, a LDA classifier was trained to discriminate between each stimulus pair at each time point, and the resulting values were placed into Representational Dissimilarity Matrices (RDMs). Intact and silhouette stimuli were analyzed separately. Next, model RDMs were created for aspect ratio and category. We focused on these two dimensions because category and aspect ratio have been proposed to explain one another in the literature (Bao et al., 2020; Yargholi & Op De Beeck, 2023). In these model RDMs, stimuli pairs that were predicted to be similar were given values of 0 (e.g. two spiky stimuli in the aspect ratio model), and stimuli that were predicted to be different were given values of 1 (e.g. one stubby and one spiky in the aspect ratio model). To determine which model best explained the representational structure of the neural data, a linear model was then conducted at each time point of the trial. Linear models were performed separately for intact and silhouette stimuli. This analysis resulted in a time course of beta values in which the beta value can be interpreted as the strength of that model’s ability to predict the neural data.

### Statistical Analysis

To test the probability of above-chance (alternative hypothesis) classification versus chance-level (null hypothesis) classification across participants at each timepoint, we conducted Bayes Factors (Teichmann et al., 2021). This analysis allowed us to directly assess and measure the strength of different hypotheses. Bayes Factors were calculated using the Bayes Factor R package (Morey et al., 2015) implemented in Matlab (Teichmann et al., 2021). Bayesian t-tests were run at each timepoint. To allow for small effects under the alternative hypothesis, the prior range of the alternative hypotheses were adjusted to be from .5 to infinity (Rouder et al., 2009). A half-Cauchy prior with the default, medium width of .707 was used (Teichmann et al., 2021). Larger Bayes Factors (BFs) indicate more evidence for the alternative hypothesis that there is above-chance classification. Specifically, BFs that are larger than 1 are evidence for above-chance decoding, whereas Bayes Factors smaller than 1 support at-chance decoding. The strength of Bayes Factors is directly interpretable, such that a BF of 5 indicates that the observed data is 5 times more likely under the alternative hypothesis compared to the null hypothesis whereas a Bayes Factor of 1/5 indicates that observed data is 5 times more likely under the null hypothesis compared to the alternative hypothesis. This Bayesian approach was implemented for the classification and temporal generalization analysis. A similar Bayesian method was also used to compare neural RDMs to model RDMs at each timepoint, however the prior range was adjusted to be two-tailed without an expected null prior value.

Additionally, Bayes Factors were used to evaluate the probability of above-chance differences in decoding, temporal generalization, and beta values between intact and silhouette stimuli for each dimension. Within each participant, we calculated the difference between decoding accuracy (for both the decoding analysis and temporal generalization analysis) and beta values for intact and silhouette stimuli at each time point and performed Bayesian t-tests against 0. We used a half-Cauchy prior with the default, medium width of .707 (Teichmann et al., 2021), and we excluded the interval between d=-0.5 and d=5 from the prior. Bayes Factors corresponding to differences between intact and silhouette stimuli are included in each figure for decoding, temporal generalization, and RSA analyses.

## Results

### Decoding analyses demonstrate unique temporal dynamics underlying representations of each object space dimension

To investigate the temporal dynamics underlying the dimensions of aspect ratio, animacy and category during object processing, we trained separate linear classifiers to discriminate aspect ratio (stubby vs spiky), category (faces vs bodies vs manmade objects vs natural objects), and animacy (animate vs inanimate). These analyses were carried out separately for intact and silhouette stimuli and resulted in a time series of decoding accuracies that revealed when information related to each dimension was decodable for each stimulus type (Figure 2).

**Figure 2.**
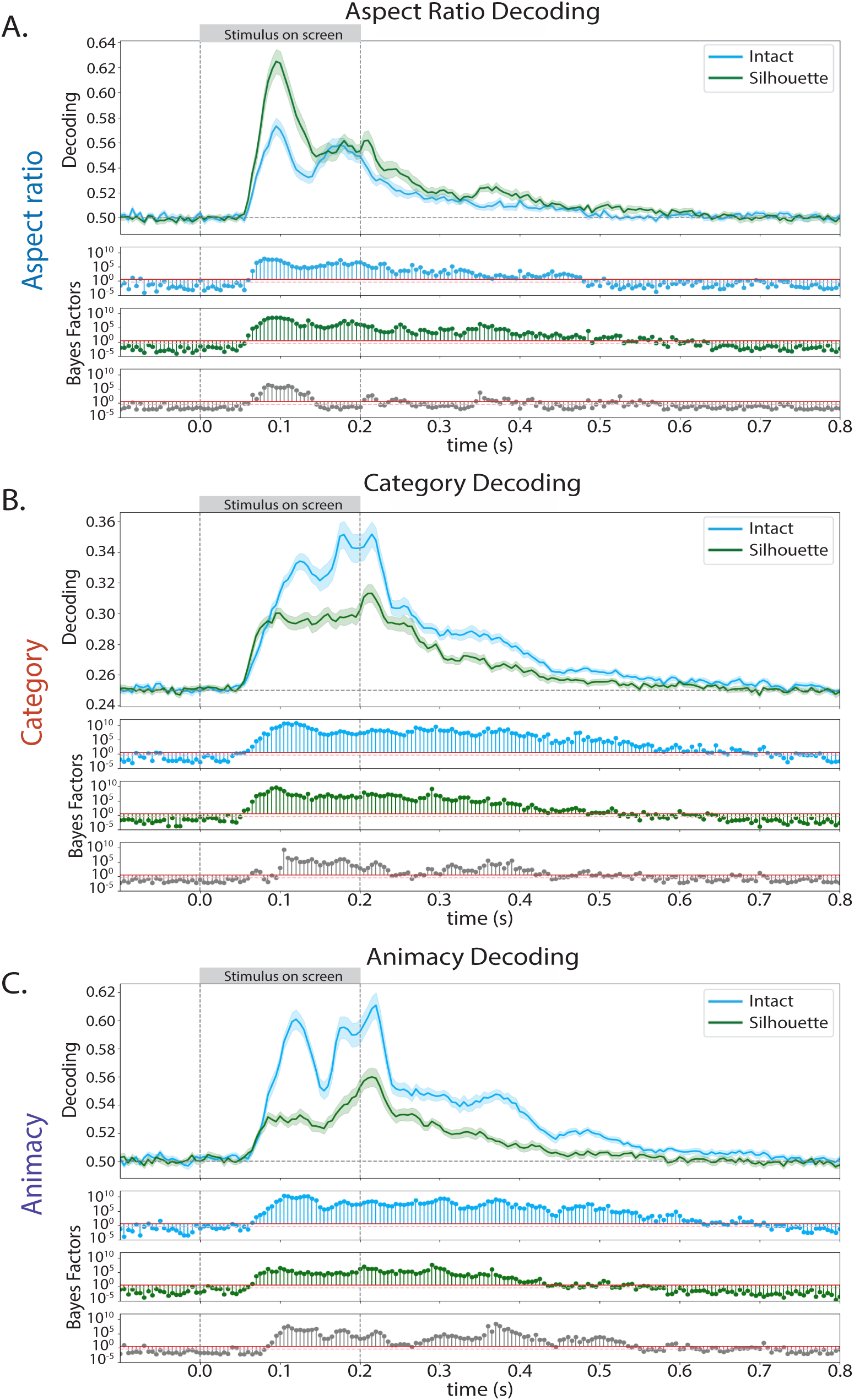
Mean decoding accuracy from whole-head EEG activation patterns for decoding. (a) the aspect ratio (stubby vs spiky) of the object, (b) the category (face, body, manmade object, natural object) of the object, and (c) the animacy (animate vs non-animate) of the object. In all plots the shaded region indicates the standard error at that time point. The corresponding Bayes Factors for each stimulus type (blue for intact stimuli and green for silhouette stimuli) are directly below the decoding accuracy plot. Bayes Factors for the difference (intact - silhouette) between decoding accuracy in the intact stimuli and silhouette versions of those stimuli and included in gray. Information about each object space dimension was available in intact and silhouette stimuli, and decoding strength of each dimension was modulated by stimulus information available.

Each dimension was decodable in both intact and silhouette stimuli, and decoding onset was similar across all dimensions for both stimulus types. In both intact and silhouette stimuli, aspect ratio decoding peaked ∼93 ms after stimulus onset, which was earlier than both animacy and category information (peak decoding at ∼118 ms and ∼125 ms, respectively). While each dimension had a second decoding peak in both intact and silhouette stimuli, this peak had lower accuracy for aspect ratio, whereas the second decoding peak was as accurate or had a higher accuracy than the initial decoding peak for both animacy and category. Above-chance decoding was sustained after stimulus offset for each dimension. In both stimulus types, aspect ratio decoding offset occurred ∼500 ms after stimulus onset, animacy decoding offset occurred ∼525 ms after stimulus offset, and category decoding offset occurred ∼550 ms after stimulus onset.

Importantly, the decodability of each dimension was modulated by the amount of available stimulus information. Aspect ratio decoding was more accurate for silhouette stimuli (peak decoding ∼63%) in comparison to intact stimuli (peak decoding ∼57%) during stimulus presentation, and strong evidence for an above-chance difference in decoding strength began before the initial decoding peak at ∼65 ms until ∼145 ms after stimulus onset (Figure 2a). This pattern was reversed for both animacy and category decoding (Figure 2b and 2c). Decoding accuracy was higher for both dimensions in intact stimuli (animacy decoding accuracy at ∼61.5% and category decoding accuracy at ∼35.5%) compared to silhouette stimuli (animacy decoding accuracy at ∼56% and category decoding accuracy at ∼31%). Evidence for above-chance differences in decoding strength occurred later (both occurring around 100 ms after stimulus onset) and for a more sustained period of time (including after the stimulus was no longer on the screen) for category and animacy when compared to the aspect ratio dimension.

These decoding results indicate that the dimensions of aspect ratio, animacy, and category have different temporal dynamics from one another. Aspect ratio information is available earlier and for a shorter time period when compared to category and animacy, and available stimulus information directly impacts the amount of information available for each dimension underlying object space.

### Temporal Generalization

To evaluate the stability of each dimension over time, temporal generalization analysis was performed. We trained and tested classifiers on different time points during the trial, resulting in a measure of information generalization during and after stimulus presentation. Off-diagonal classification indicates information is transiently represented over those time points, whereas classification that looks like a square near the diagonal, or classification found in the upper or lower triangle of the graph suggests information stability across time.

Once again, different temporal dynamics were observed for aspect ratio in comparison to animacy and category (Figure 3). Almost no generalization was found for aspect ratio in intact stimuli, and no above-chance classification occurred after 500 ms post stimulus onset (Figure 3a). In silhouette stimuli, above-chance generalization occurred beginning ∼86 ms after stimulus onset, and remained after stimulus offset. Additionally, there was off-diagonal generalization, in which early time points between 86 ms and 185 ms post stimulus onset generalized through 600 ms after stimulus offset (Figure 3b). This pattern indicates that aspect ratio information is more transiently available during visual processing of the intact stimuli, and for silhouette object stimuli, aspect ratio information is held consistently over time. In contrast, generalization of animacy and category information over time were found during object processing of the intact stimuli (Figure 3a). Above-chance animacy classification began around 90 ms after stimulus onset, and again around 180 ms after stimulus onset through to 35 ms after stimulus offset. Small, but above-chance, generalization also occurred between ∼300 ms and ∼550 ms after stimulus offset. Off-diagonal classification was found between the timepoints of ∼200 ms and ∼550 ms after onset. Category classification had almost the exact same pattern of generalization, and off-diagonal decoding was found for multiple periods of time during and after stimulus presentation in the intact stimuli data (Figure 3b). There was almost no evidence for above-chance off-diagonal classification, apart from weak classification between ∼480 ms to 600 ms after stimulus offset. These results re-enforce that aspect ratio has differential temporal dynamics from category and animacy. Additionally, these data support that amount of stimulus information available does not only impact when object space dimensions are represented in the neural signal, but also the stability of that representation over time. For all dimensions, evidence for above-chance differences between the temporal generalization accuracies in the intact stimuli and the silhouette were observed (Figure 3c).

**Figure 3.**
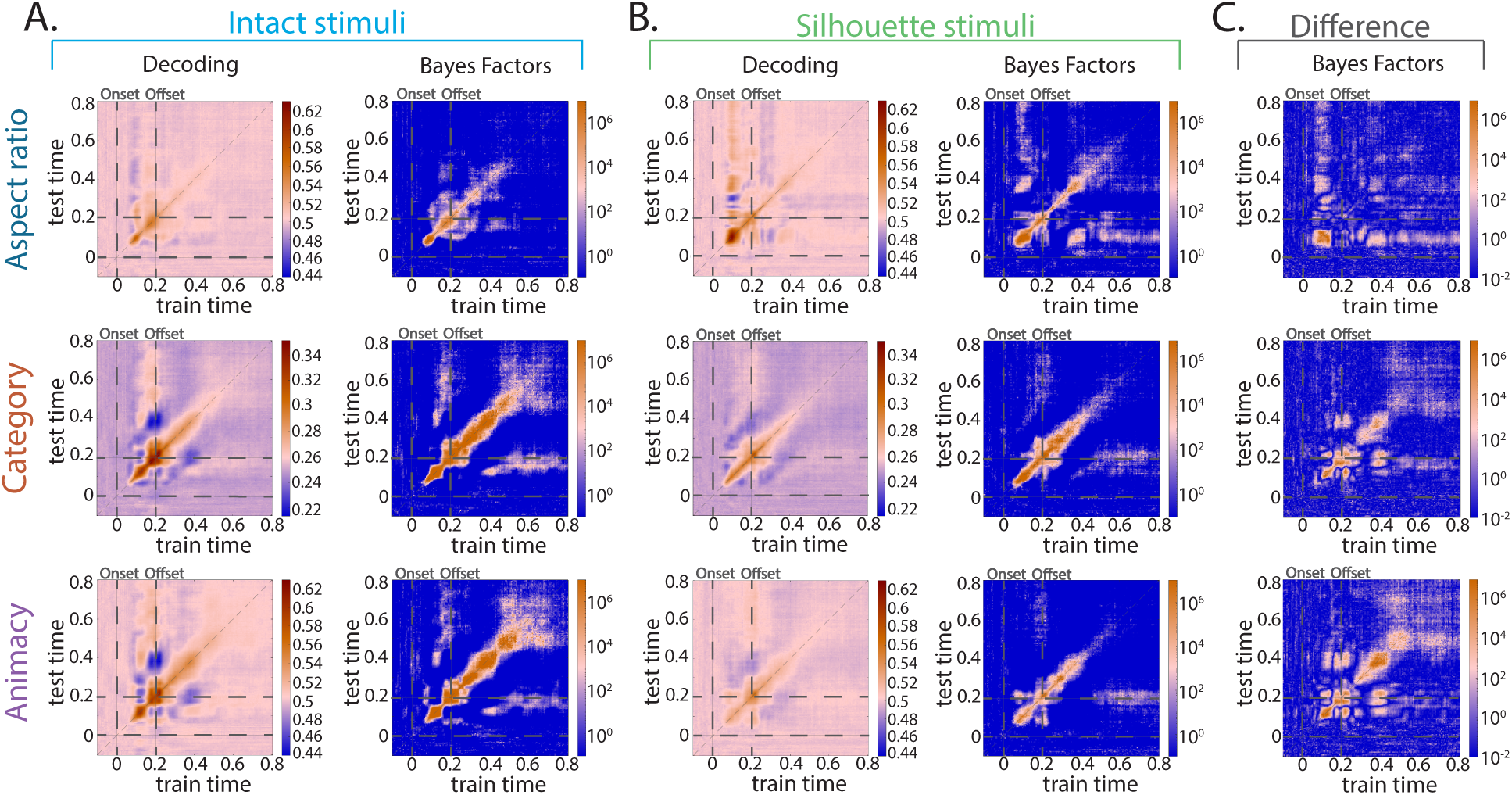
Mean temporal generalization for decoding each dimension in (a) the intact stimuli, and in (b) silhouette stimuli. For all decoding plots, pink indicates chance-level temporal generalization decoding accuracy values, while warm orange-red colors indicate above-chance decoding accuracy values. (c) The Bayes Factors corresponding to the difference in temporal generalization decoding accuracy for each dimension between intact and intact stimuli. For all Bayes Factors plots, warm colors (pink and light-to-dark orange) indicate Bayes Factors above 1 (evidence for the alternative hypothesis), while blue indicates Bayes Factors below 1 (evidence for the null hypothesis). The points along the diagonal of each plot represent testing and training on the same time point. In the intact stimuli, stability in category and animacy information was observed, while in the silhouette stimuli stability in aspect ratio information was observed. Strong evidence for above-chance differences between temporal generalization in the intact stimuli and the silhouette stimuli for each dimension was found.

Overall, these results indicate that aspect ratio information is more transiently represented when compared to category and animacy information in stimuli in which internal details are preserved. However, this pattern is reversed in the silhouette stimuli, mirroring the decoding results. In stimuli without internal details, aspect ratio information is more strongly represented and is more stable when compared to animacy and category information.

### Representational Similarity Analysis

Lastly, to examine which object space dimension best explains the representational structure of the neural data, we performed a linear model between the neural representational dissimilarity matrix and model dissimilarity matrices representing aspect ratio and category at each time point (Figure 4). This resulted in a time course of beta values at every time point, which can be interpreted as the relative strength that model explains the observed neural data, while taking into the other model into consideration. We chose to focus on aspect ratio and category in this analysis, because conflicting results were found for these dimensions in previous proposed maps of object space (Bao et al., 2020; Coggan & Tong, 2023; Yargholi & Op De Beeck, 2023). We also performed this analysis when including a model representing animacy and the results were largely preserved as they are reported here.

**Figure 4.**
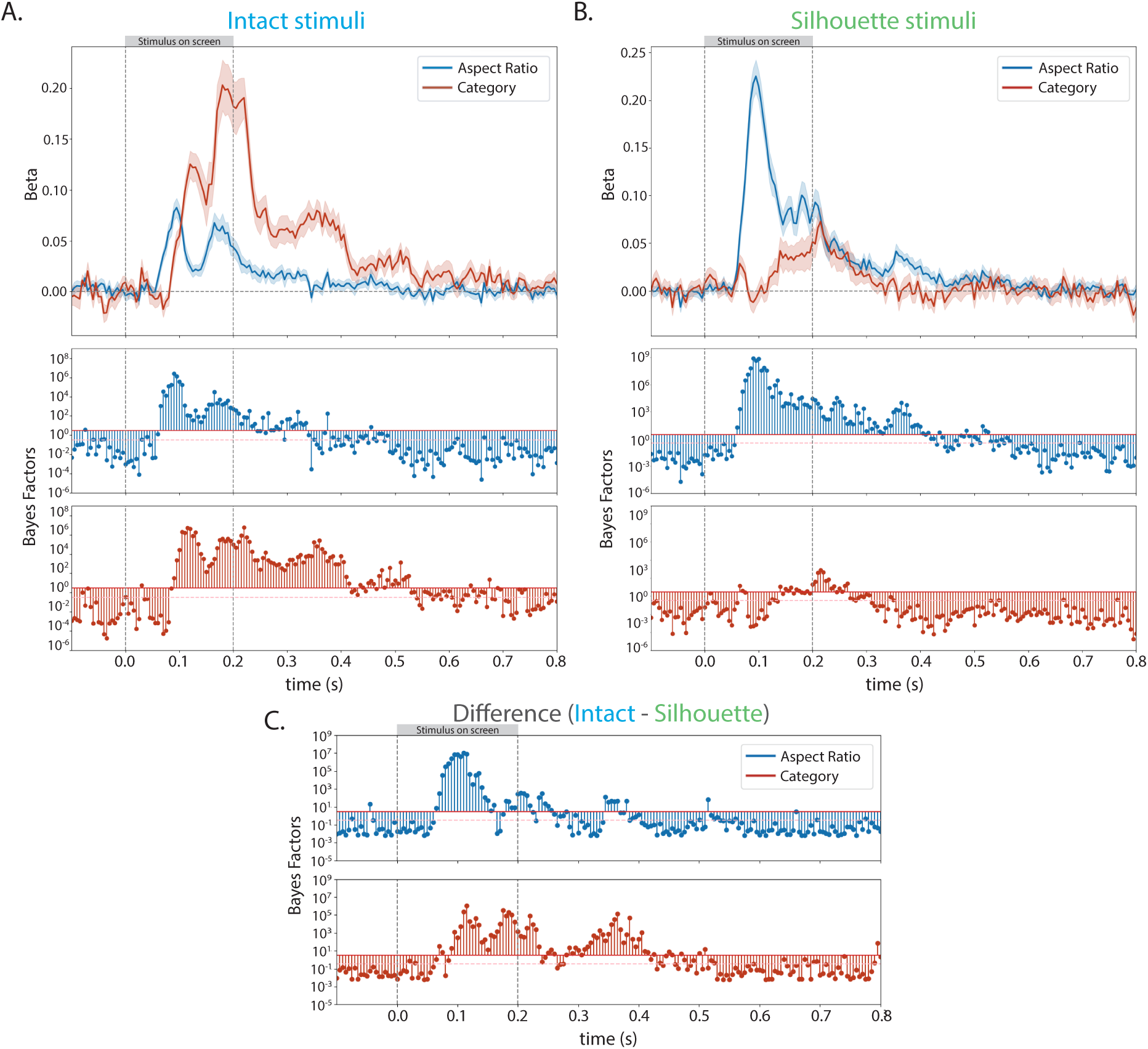
Mean beta values corresponding to models of aspect ratio and category calculated via linear models in (a) intact stimuli and, (b) silhouette stimuli. In all plots the shaded region indicates the standard error at that time point, and the corresponding Bayes Factors for each stimulus type (blue for intact stimuli and green for silhouette stimuli) are directly below the betas plot. (c) Bayes Factors for the difference (intact - silhouette) between beta values in the intact stimuli and silhouette versions of those stimuli during the entire trial. The category model better explains the representational structure of the neural data for the intact stimuli, and the aspect ratio model better explains the representational structure of the neural data for the silhouette stimuli.

The aspect ratio model explained the neural data best for both intact and silhouette stimuli beginning around 70 ms and peaked around 95 ms after stimulus onset (Figure 4). For intact stimuli (Figure 4a), the category model then best explained the neural data beginning around 105 ms after stimulus onset for the rest of the time the stimulus was on the screen until ∼225 ms after stimulus offset. The aspect ratio model uniquely explained the neural data throughout the trial as well until approximately 20 ms after stimulus offset. In contrast, there was no evidence that the category model explained the neural data for silhouette stimuli at almost any point in the trial (briefly for ∼25 ms after stimulus offset) (Figure 4b). Instead, aspect ratio information best explained the neural signal for the entirety of the time the stimulus was on the screen, and for 200 ms after stimulus offset. These data provide evidence that when object details are concealed, aspect ratio information better explains the neural signal during visual and post-visual processing.

Bayes Factors showed strong evidence for differences between the betas in the intact stimuli and silhouette stimuli (Figure 4c) for the aspect ratio and category models. These differences primarily occurred earlier for the aspect ratio model and occurred more consistently later in the trial, including after stimulus offset, for the category model. These data suggest that both aspect ratio and category information explain unique components of the neural signal when viewing intact and silhouette objects. Category information better explains the neural data for a longer period of time in the intact stimuli, however, does not explain the neural data at almost any point in the silhouette stimuli.

## Discussion

The goal of this experiment was to evaluate if, and when, representations of proposed object space dimensions were available during visual processing of intact object stimuli and object stimuli with masked internal details. Specifically, we investigated the temporal dynamics of aspect ratio, category, and animacy using multivariate pattern analysis of whole-head EEG data activation patterns. We found that representations of each dimension were available in the neural signal in both stimulus types. Aspect ratio information was available earlier than category and animacy information, and the strength of decoding was modulated by stimulus information available during visual processing. Aspect ratio information was also more transiently available when compared to category and animacy information in the intact stimuli, both of which were stable over time. However, in silhouette stimuli, aspect ratio information was more stable than category and animacy information. Models of aspect ratio and category object information differentially explained the representational structure of the neural data over time in the intact stimuli, and aspect ratio best explained the neural data in the silhouette stimuli during and after stimulus presentation. Overall, these results demonstrate that each dimension of object space has distinct temporal dynamics, and the amount of stimulus information available during visual object processing directly modulates the representations of these dimensions over time. Importantly, our findings highlight the importance of considering temporal information when defining object space (Robinson et al., 2023).

The fundamental organizing principles of object space and higher order visual regions have been the subject of several studies and debates. For example, a model of object space in macaque inferotemporal cortex was recently proposed, defined by orthogonal dimensions of aspect ratio and animacy (Bao et al., 2020). This model accounted for category information (e.g. face and body representations) as a result of the four quadrants of this two-dimensional object space. Some evidence for these organizing principles were found in human inferotemporal cortex, however a clear sequence of this object space was not observed, and the selectivity of aspect ratio-animacy space highly overlapped with category selectivity (Coggan & Tong, 2023). When using a stimulus set that dissociated category and aspect ratio information, object space and the organization of human occipitotemporal cortex was better characterized by the dimensions of category and animacy. In this model, aspect ratio was not represented, and this dimension was instead explained by category effects (Yargholi & Op de Beeck, 2023). These diverging conclusions could be a result of the different stimulus set, different amounts of category information available between species (e.g. object category familiarity), or the lack of temporal sensitivity in human fMRI compared to macaque electrophysiology data. Our results directly speak to these possible explanations. First, we used the stimulus set that dissociated aspect ratio and category information and created a silhouette version of those stimuli, effectively masking internal stimulus details and reducing available category information. Secondly, we used EEG to evaluate changes over fine-grained periods of time during object processing. We demonstrated that category and aspect ratio have distinct and differentiable temporal dynamics, both of which uniquely explain the neural data at different time points. Representations of these dimensions were modulated over time by binarizing the stimuli. Therefore, it is possible that the stimulus set confounding category and aspect ratio, combined with a lack of available category information (Bao et al., 2020) resulted in aspect ratio explaining category information in the macaque inferotemporal cortex data, while the available category information combined with the slower resolution of fMRI resulted in category information explaining the dimension of aspect ratio in human participants (Coggan & Tong, 2023; Yargholi & Op de Beeck, 2023). Overall, our results indicate that the dimensions of aspect ratio and category are both represented during object processing and one object space dimension does not fully explain the other dimension. Our data suggest that object space is flexible, and likely weights dimensions underlying object space differently depending on the visual information that is available.

Our results corroborate studies that have investigated the dissociation of visual features, such as aspect ratio and shape, from category information during object processing. While shape and aspect ratio are distinguishable from one another (Bougou et al., 2024; Bracci & Op De Beeck, 2016) and shape is an overall coarser and less standardized visual feature, they are highly related and will both be discussed here. In single unit recordings in macaque inferotemporal cortex, visual features such as aspect ratio (along a gradient of round to thin shapes) better explained neural responses when compared to semantic or category information (Baldassi et al., 2013). Dissociations between category information and both aspect ratio and shape were also reported in human multi-unit activity and high-gamma responses in the Lateral Occipital Complex (Bougou et al., 2024). Similar findings have been reported when investigating shape. Within body stimuli, shape information was preserved independent of category information, and this shape information was available earlier than category information (Kaiser, Azzalini, et al., 2016). Several fMRI (Bracci & Op De Beeck, 2016; Kriegeskorte et al., 2008; Op De Beeck et al., 2008; Proklova et al., 2016) and MEG (Carlson et al., 2013; Cichy et al., 2014, 2016; Contini et al., 2017) studies have also observed shape information independent of category information and vice versa during object processing. While not all of these studies specifically dissociated category from shape within the presented stimuli, post-hoc analysis accounting for visual properties found shape and category could not fully explain one another. Interestingly, for unfamiliar objects, the organization of human object-selective regions were strongly related to perceived shape information (Op De Beeck et al., 2008), supporting the idea that aspect ratio and shape information may be more strongly weighted when category and object identity information is not available. Our results extend this literature by exploring the temporal dynamics of stimuli where category and aspect ratio are orthogonalized, and by masking internal object details in these same stimuli. Aspect ratio information is stronger and more stable over time in silhouette objects in comparison to intact objects and is available earlier than category information in all objects. Future studies may benefit from explicitly investigating the relationship between object discernability and the temporal dynamics of object space dimensions, and by investigating whether this directly translates into performance on behavioral tasks that prioritize shape versus category (Grootswagers et al., 2018; Koenig-Robert et al., 2024).

Recently, a new approach to investigating object processing has been proposed (Contier et al., 2024). When compared to traditional category or feature-driven approaches, object space dimensions (66 in total) derived from millions of behavioral responses could better explain neural responses across human visual cortex. These dimensions also replicated and explained previously-reported feature and category selectivity (Contier et al., 2024). When investigating the temporal dynamics underlying these behaviorally-relevant object space dimensions, multidimensional information was rapidly available (∼80 ms) and physical object property information was available earlier than conceptual information (Teichmann et al., 2023). These results highlight that object space is dynamic over time and that object processing needs to be considered more holistically than low-dimensional object spaces. Specifically, object processing is likely a result of a complex interplay between external stimulus information (Grootswagers, Robinson, & Carlson, 2019), co-occurring object dimensions (Guntupalli et al., 2016; Haxby et al., 2011) that are relevant for behavior (Contier et al., 2024; Grootswagers et al., 2018; Teichmann et al., 2023) or social interaction (Contini et al., 2020; Grootswagers et al., 2022b), and are sensitive to task demands (Grootswagers et al., 2021; Kaiser, Oosterhof, et al., 2016) and individual differences (e.g. internal representations, cognitive abilities) (Bracci & Op De Beeck, 2023). Our results can be easily interpreted through this multidimensional framework, such that the object space dimensions we probed were distinct from one another even though they co-occurred, dynamically evolved over time, and were flexibly weighted depending on available stimulus information. In future studies, it would be helpful to investigate the temporal dynamics underlying behaviorally-derived dimensions in both intact and silhouette stimuli across varying task demands to more fully characterize object processing and understand the importance of object space dimensions in different conditions over time.

In sum, the object space dimensions of aspect ratio, category, and animacy are all present and distinct from one another over time and differentially explain the representational structure of the neural data during object processing. The strength and stability of these dimensions are modulated by the absence of internal object details, suggesting that the object space is flexible and depends on the information available to the visual system. These results help resolve previously reported discrepancies in the literature and support a more multidimensional view of object space. Moreover, these data reinforce that temporal information is important to consider when investigating object processing. Object processing, and therefore object space, is not static and evolves over time, and should be investigated using methods that are sensitive to these dynamics.

## Data and Code Availability

Code is available on GitHub (https://github.com/alexiskidder/ObjectProcessing_RSVP) and data is available on OpenNeuro.

## Author Contributions

**Alexis Kidder**: Conceptualization, Methodology, Software, Formal Analysis, Investigation, Data Curation, Writing – Original Draft, Writing – Review & Editing, Visualization

**Genevieve L. Quek**: Conceptualization, Validation, Resources, Writing – Review & Editing, Supervision

**Tijl Grootswagers**: Conceptualization, Methodology, Software, Validation, Resources, Writing – Review & Editing, Supervision

## Funding

This work was supported by the International Visiting Scholar Program at The MARCS Institute for Brain, Behaviour, and Development, Western Sydney University and by the Australian Research Council grant DE230100380.

## Declaration of Competing Interests

The authors have declared that there are no conflicts of interest in relation to the subject of this study.

## Acknowledgements

The authors would like to thank Lina Teichmann and Chris Baker for helpful discussions, and Mahdiyeh Khanbagi, Jessica Chen, Nanzin Sheykh Andalibi, and Almudena Ramírez Haro for assistance with data acquisition and EEG methods.

## Notes

### Competing Interest Statement

The authors have declared no competing interest.

